# Effects of human-wildlife Conflicts on Local People’s Livelihoods and Wildlife Conservation in and around Alitash National Park, Northwest Ethiopia

**DOI:** 10.1101/2022.08.29.505656

**Authors:** Mekuriaw Zewdie Ayalew, Getahun Tassew Melese

## Abstract

Human-wildlife conflict becomes one of the fundamental aspects of wildlife management because it threatens both wildlife and human. People injured, abused, and killed wildlife in response to the perceived and/or actual damages from wildlife. Such negative interaction between humans and wildlife exists everywhere but more intense and costly in adjacent protected areas. This study aimed to assess the effects of human-wildlife conflicts on local people’s livelihood and wildlife conservation in and around Alitash National Park. The primary data were collected using multi-stage sampling techniques by employing a combination of social survey methods involving participatory techniques (focus group discussions and key informant interviews) and structured household interviewers. The survey questionnaire consisted of both open-ended and fixed-response questions designed to solicit information mainly for wildlife-caused damages and other costs to the local community along with human-induced wildlife decline and perceptions towards wildlife conservation. The results revealed that 59.6% of the respondents encountered at least a single type of damage. More than half (54%) of respondents encountered crop damage, of which 34.4% lost up to 25% of their crop fields whereas the amount of loss to the remaining (18.7%) was to 75% of the crop fields. The livestock loss during the last five years of the period prior to the survey administration was 287.89 tropical livestock units which were shared by 47.5% of the respondents. Wildlife attacks on humans were rare but eighteen attacks were caused by Panther leo, Crocuta crocuta and Panthera pardus were encountered when the victims cross the habitats of the species. The responses of the victims to the wildlife imposed the damage was negative and full of anger and resentment. This has been undermining conservation efforts and the socio-economic welfare of the local people around Alitash National Park. Providing alternative livelihood opportunities and creating a context-based conservation scheme along with continuous conservation education would help to reduce the negative effect of human-wildlife conflicts on both wildlife and people in the study area.

## Introduction

Human-wildlife conflict occurs when the needs and activities of wildlife undermine human welfare, health, and safety and/or when human negatively affect the need and survival of wildlife (Madden, 2004). The conflict threats both human livelihood and wildlife conservation (Ladan, 2014) as wildlife cause damage to livestock, crop, property and man lives while human beings persecute wildlife in retaliation for losses incurred, and undermine their conservation and survival through poaching, illegal hunting and destruction of their habitats (Ogra, 2008). Such negative interactions between humans and wildlife become the most complex challenge to wildlife conservation and the socioeconomic welfare of rural households around the world (Eniang et al., 2011).

Although human-wildlife conflict is a global concern, it is more serious in developing countries than developed nations due to the reason that agriculture is the mainstay of the local livelihood (Lamarque et al., 2009). Most rural communities in Ethiopia and other developing countries depend on mixed farming (crop cultivation and livestock rearing) for their source of food and income (Ladan, 2014). Therefore, any damage on crop and livestock greatly jeopardizes their socioeconomic well-being (Lewa et al., 2017). The damages affect the farmers’ ways of life and ability to feed their families (Eniang et al., 2011). Missing school, work, social events and sleep to guard crop and livestock day and night from wildlife are other costs (Mojo et al., 2014) to farmers adjacent protected areas. In Ethiopia, crop and livestock loses are a significant for farmers living closer to wildlife areas (Haylegebriel, 2016; Leta et al., 2016; Teshome & Girma, 2017) although the costs in health and social settings are also undeniable (Lamarque et al., 2009).

Most protected areas in Ethiopia become islands surrounded by seas of cultivation and settlement (Alemu et al., 2017) which causes serious pressure on the protected areas and results serious habitat loss. Following the pressure and the habitat loss, wild animals stray to nearby grazing, farming and settlement areas in search of their requirements (Madden, 2008). In doing so, wildlife impose intolerable economic, social and health risks to the local people and hence people retaliate against conservation authorities by killing wild animals (Biset et al., 2019), taking resources under protection and facilitating outside poachers (Leta et al., 2016). These antagonistic behaviors towards wildlife (Haylegebriel, 2016) have been growing among the local people living around protected areas of Ethiopia (Teshome & Girma, 2017) which generally results failure in wildlife conservation efforts (Eniang et al., 2011).

Regardless of increment in number of protected areas in Ethiopia, wild animal population has been declining (Berhanu & Teshome, 2018). Though there exist more than 55 protected areas, about 17.1% of the total area of Ethiopia, (Mamo, 2017), most of them are in poor status and their wildlife is declining (Mengist, 2020). Considering the actual population growth rate adjacent to protected areas and the growing demand for natural resources, the conflict between human and wildlife is inevitably intensified (Ladan, 2014) as it is the case in Alitash National Park (Agitew et al., 2016). To this end, bringing remarkable balance between conservation priorities and the need and burden of communities residing adjacent protected areas is highly crucial (Teshome & Girma, 2017). However, it requires a comprehensive understanding about consequences of human-wildlife conflicts on both wildlife and people but yet untapped by previous studies. This study, therefore, aimed to assess effects of human-wildlife conflicts on both people and wildlife in and around Alitash National Park.

## Materials and Methods

### Description of the Study Area

Alitash National Park, the study area, upgraded from priority forest area to National Park in 2006 with the regional legal issues publication newsletter ‘Zekere heg’ by Amhara Regional Council Regulation Act No 38/2005 (PaDPA, 2009). The name Alatish was used to designate the park but latter it has been changed into Alitash by the Councils of Ministers Regulation No.333/2014. Alitash National Park is located in West Gondar administrative zone particularly at Quara district between 110 47’ - 120 21’ N latitude and 350 16’ - 350 47’ E longitude (Fig.1). It is far about 1,123 km from Addis Ababa (the capital of Ethiopia) and 534 km from Bahirdar (the capital of Amhara National Regional State) (PaDPA, 2009). Covering 2665 km^2^ area of land, the Park shares boundaries with Sudan (Dinder National Park) in the west, Benshangul Gumuz National Regional State in the south and different peasant associations of Quara district in the eastern and northern edge (Agitew et al., 2016).

**Figure 1.**
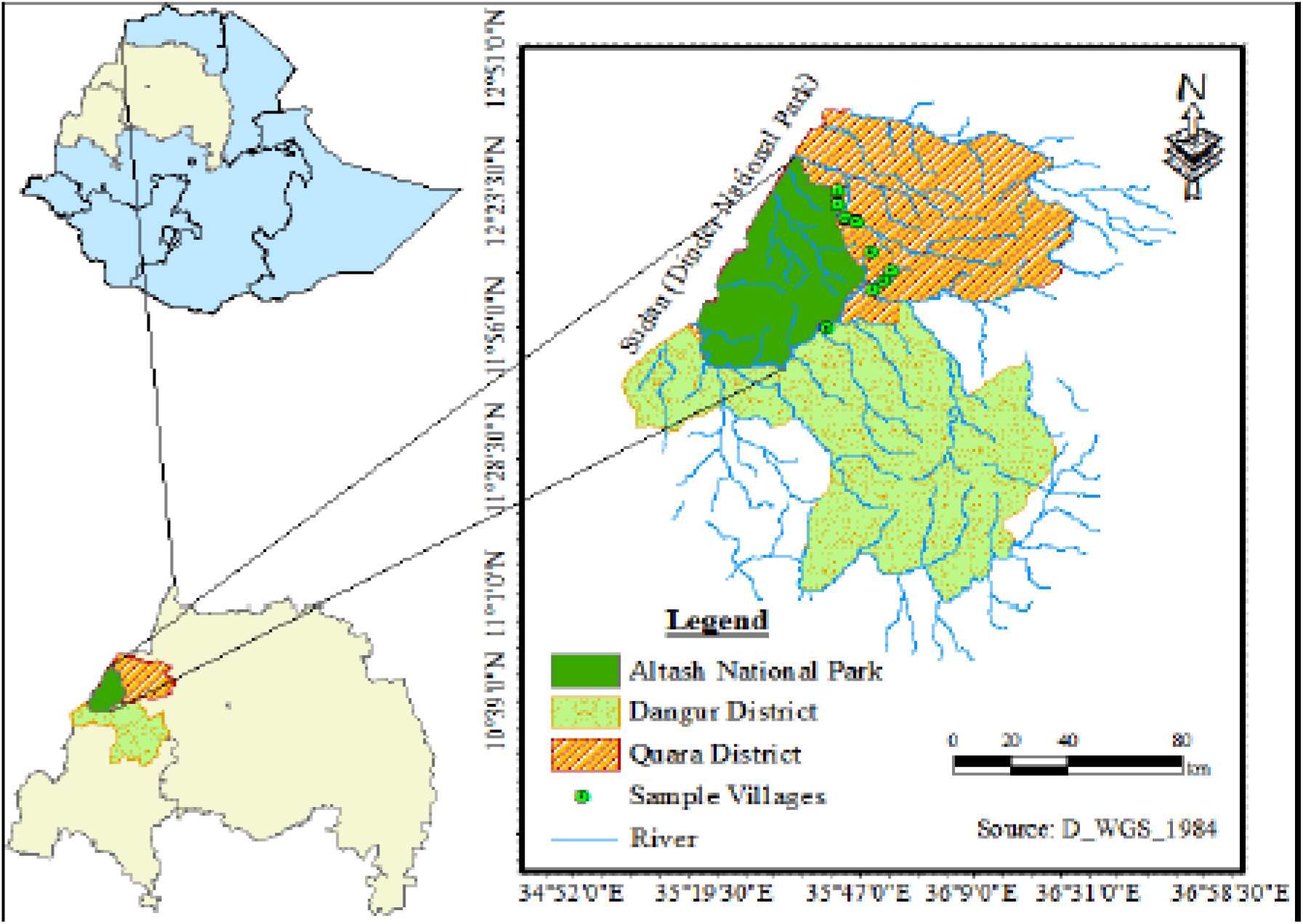
Map of the study area

Alitash National Park is rich in floral and faunal diversity. The Park consists of natural vegetation composed of 176 species of plants of which 29 herbaceous and 17 grasses that can be used as a shelter and feed sources for the wildlife hosted in it. It hosts 37 mammalian, 204 bird and 23 rodents species as well as 6 insectivores and 7 types of reptiles and amphibians (PaDPA, 2009). Among the mammalians, there are endangered and rare species like African elephant (*Loxodonata africana*), Leopared (*Panthera pardus*), lion (*Panthera leo*) and also the lower risk but conservation dependent species (*Tragelaphus imberbis* and *Tragelaphus strepsiceros*) are the integral part of the park biodiversity (Ayalew & Melese, 2020).

### Sampling Design

The study employed multi-stage sampling procedure. All peasant association adjacent to Alitash National Park was purposively selected in the first stage for the presence of conflicts between human and wildlife. The villages of the peasant associations adjacent to the park were the focus area of the study by assuming that people in close proximity to wildlife areas might tend to have more affected by human-wildlife conflicts (Angela et al., 2014) and are more aware about the problem than those far (Nuno et al., 2013).

In the second stage of the sampling, the villages were stratified based on their proximity to the park as nearest (less than 7km), medium (7-14km) and far (greater than 14km) (Bezihalem et al., 2017). Among the villages in the three strata, eleven villages were selected using proportional random sampling technique. In the third stage, sample households were selected randomly in each village. To determine the sample size, the formula developed by Cochran (1963) was employed. Accordingly, one hundred ninety eight (198) households were selected proportionately.

### Data Sources and Collection

Both primary and secondary data were collected for this study. The primary data were collected by employing a combination of social survey methods involving participatory techniques (focal group discussions and key informant interviews) and structured household surveys. Three focus group discussions comprising 7-12 diversified discussants and key informant interviews with 12 key informants were conducted to consolidate and triangulate the data collected through the household surveys. The household survey questionnaire consisted of both open-ended and fixed-response questions designed to solicit information that mainly focus on wildlife caused damages and other costs to the local communities, perception and antagonistic behavior of local people towards wildlife and wildlife conservation. Moreover, a series of supplementary questions were included in the questionnaire to gather respondents’ socio-economic and demographic information. The interviews were mainly target on the head of households but in case of their absence other permanent resident adult (≥18 years) member of a household were allowed to took part in the interview. In absence of a suitable respondent, the adjacent household was surveyed instead. The interviews were conducted in face to face fashion at each respondent’s homestead by well-trained interviewers.

### Methods of Data Analysis

Based on the objectives of the study and nature of the data collected, different data analysis techniques were employed using the statistical package for social science (SPSS) version 20 throughout the whole analysis process. To analyze the collected data both descriptive and inferential statistics were applied. A chi-square (χ2) test for goodness-of-fit was used to establish whether or not the sampled households’ socio-demographic composition data are significantly differed. In addition, cross tabulations involving chi-square (*χ*2) tests was employed first, to establish whether responses on prevalence and trends in human-wildlife conflicts are dependent and/or independent of location within the study villages, and second, to establish the relationships between socio-demographic variables and households’ responses on prevalence and perceived trends of human-wildlife conflict. Binary logistic regression analysis was applied to identify the major wildlife caused damages in shaping the local people’s perception towards wildlife and their conservation. All tests were considered significant at *P* ≤ 0.05.

## Results

### Socio-demographic Characteristics

The data were collected from 198 households. Among the surveyed households, 34.4% were dwelling within the range of 7 km distance from Alitash National Park boundary whereas 39.3% were residing 7 to 14 km far from the park. The remaining (26.7%) were within the range of 14-21 km distance from the edge of the National Park.

Most of the respondents (96%) of the study were male and their age distribution varied from 18 to 71 years. A little more than half (51%) of them were from 36 to 50 years of age and the rest, 18 to 35 and 51 to 71 years of age covered 26.3% and 22.7%, respectively. About 47% of the respondents were uneducated, 37.9% of them were able to read and write whereas some (9.6%) joined primary school and the rest (5.6%) attended at least high school education.

The finding revealed that most (86.9%) of the surveyed households practiced the combination of crop and livestock farming for their livelihood while 11.1% of them used crop cultivation or livestock rearing together with off-farm activities like trading, illegal hunting, charcoal making and as traditional healers. The remaining (2%) were tied to the off-farm activities.

### Types and prevalence of wildlife Caused Damages

The majority (59.6%) of the respondents reported that there were crop damages, livestock predation and human attack due to the action of wildlife. Crop damages and livestock predation were the most frequently mentioned problems facing the surveyed households in the study area. Crop damage was the most prevalent (53.5%) damage type while livestock predation was the second most rated (47.5%) problem. Although wildlife attack on human was the least (8.6%) problem, it was perceived as a very serious and intolerable to the local people (Table 1).

**Table 1:** Proportion of Respondents faced wildlife caused damages

| Villages | n | Proportion of Respondents Faced wildlife caused problems (%) |  |  |  |
| --- | --- | --- | --- | --- | --- |
|  |  | Crop damage | Livestock predation | Human attack | All damages |
| Shimelegir | 15 | 73.3 | 73.3 | 0 | 73.3 |
| Bermil | 16 | 36.4 | 36.4 | 27 | 40.9 |
| Bambahoo | 14 | 80.0 | 80.0 | 6.7 | 86.7 |
| Aygelie | 13 | 70.6 | 64.7 | 23.5 | 70.6 |
| Baywa | 11 | 42.9 | 57.1 | 21.4 | 71.4 |
| Workit | 26 | 65.4 | 53.8 | 0 | 73.1 |
| Derehasen | 20 | 65.0 | 35.0 | 0 | 70.0 |
| Marwuha | 9 | 70.0 | 60.0 | 10 | 70.0 |
| Mahadid | 31 | 40.6 | 28.1 | 3.1 | 40.6 |
| Dizagumz | 16 | 25.0 | 18.8 | 0 | 25.0 |
| Mirtgelegu | 10 | 27.3 | 45.5 | 9.1 | 54.5 |
| Overall | 198 | 53.5 | 47.5 | 8.6 | 59.6 |

Prevalence of crop damages was varied among villages with different proximity to Alitash National Park. Unexpectedly, the highest prevalence (24.2%) of the problems were reported from the villages far 7 to 14Km from the park boundary whereas the least (13.6%) was in villages located within 7km range from the park boundary. The reason might be the level of vigilance among the farmers in villages closer and distant to the park boundary. Farmers living closer to the park are expected to be more vigilant than those at distant villages and thus able to prevent wildlife caused damages.

Likewise, livestock damage was more prevalent in villages located in the range of 7 to 14 km distance from the park boundary than those located nearest to the boundary. The reason might be the intensity of using the park and its surrounding among the households living at different proximity with the boundary of the park. Households at the farthest away from the park boundary use the park and its buffer zone as grazing area to their livestock and spend the night at their temporal byre built there. Since the domestic animals are within the range of wildlife habitats day and night, they are more vulnerable to predation. Certainly, households living closer to the park also use the same area for grazing but they overnight at home by moving their livestock on daily basis. For that reason, the predation event they encountered is less as compared with those overnight within the wildlife area with their livestock. However, attack on human was relatively higher in closer villages. Among the respondents in these villages, 5.1% were encountered wildlife attacks ranging from simple injury to death whereas the least record was in the distant villages.

About 41% of the respondents reported both crop damage and livestock predation, while some (12.1%) reported only crop damage and a few (6.1%) faced only livestock predation. These were consistent with the key informants’ view. However, the corresponding figures were significantly differed across the study villages (χ ^2^ = 72.69; df = 30;p = 0.000). Most respondents from Shimelegir (73.3%), Bambahoo (73.3%), Aygelie (64.7%) and Marwuha (60%) encountered both crop and livestock depredation (Table 2). The double lose happened because of shared boundary between grazing area, crop fields and wildlife habitats in the study area.

**Table 2:** Proportion of respondents encountered both crop and livestock depredation

| Villages | Proportion of Respondents Faced Damage (%) | | | df | $\chi^2$ | P-value |
| --- | --- | --- | --- | --- | --- | --- |
|  | Only crop (%) | Only livestock (%) | Both (%) |  |  |  |
| Shimelegir | 0 | 0 | 73.3 | 30 | 72.69 | 0.000 |
| Bermil | 4.5 | 4.5 | 31.8 |  |  |  |
| Bambahoo | 6.7 | 6.7 | 73.3 |  |  |  |
| Aygelie | 5.9 | 0 | 64.7 |  |  |  |
| Baywa | 14.3 | 28.6 | 28.6 |  |  |  |
| Workit | 19.2 | 7.7 | 46.2 |  |  |  |
| Derehasen | 35.0 | 5.0 | 30.0 |  |  |  |
| Marwuha | 10.0 | 0 | 60.0 |  |  |  |
| Mahadid | 12.5 | 0 | 28.1 |  |  |  |
| Diza-gumz | 6.2 | 0 | 18.8 |  |  |  |
| Mirtgelegu | 9.1 | 27.3 | 18.2 |  |  |  |
| Average | 12.1 | 6.1 | 41.4 |  |  |  |

According to the focus group discussants and key informants, the top ranked wild animal species responsible for crop damage were common warthog (*Phacochoerus africanus*), African porcupines (*Hystrix cristata*), common baboon (*Papio cynocephalus Anubis*), vervet monkey (*Ceropithecus pygerythrus*) and patas monkey (*Erythrocebus patas*) among others. However, most respondents (82.3%) agreed that crop loss to common warthog was a severe elsewhere the study villages followed by common baboon and porcupine (Table 3).

**Table 3:** Major wild animals causing crop damage as ranked by respondents

| Name of problem animal |  | Damage level as ranked by respondents (%) |  |  |
| --- | --- | --- | --- | --- |
| Common | Scientific | Major | Minor | No damage |
| warthog | *Phacochoerus africanus* | 82.3 | 13.1 | 4.5 |
| Baboon | *Papio anubis* | 74.7 | 17.2 | 8.1 |
| Porcupine | *Hystrix cristata* | 61.1 | 28.8 | 10.1 |
| Vervet monkey | *Ceropithecus pygerythrus* | 44.9 | 41.4 | 13.6 |
| Patas monkey | *Erythrocebus patas* | 38.9 | 49 | 12.1 |
| Mongoose | *Herpestes sanguineus* | 26.8 | 59.1 | 14.1 |

### Perceived Trends of Wildlife Caused Damages

Perceptions on the trend of wildlife caused damages varied among the respondents of the study (*χ*^2^ = 46.03; df = 2; p = 0.000). The higher proportion (51%) of the respondents perceived that wildlife caused damages had increased whereas 36.9% perceived that the damages had decreased, and a few (12.1%) perceived remained the same since 2005. The remaining responses were mixed.

Increment in trend of crop damages by wildlife were reported by majority of respondents from Marwuha (100%), Shimel egir (86.7%), Bambahoo (66.7%), Derehasen (65%), Bermil (54.5%) and Workit (53.8%). Conversely, most respondents from Mahadid (81.2%), Mirt-gelgu (81.8%) and Baywa (57.1%) perceived the decreasing trend of such damages (Table 4).

**Table 4:**
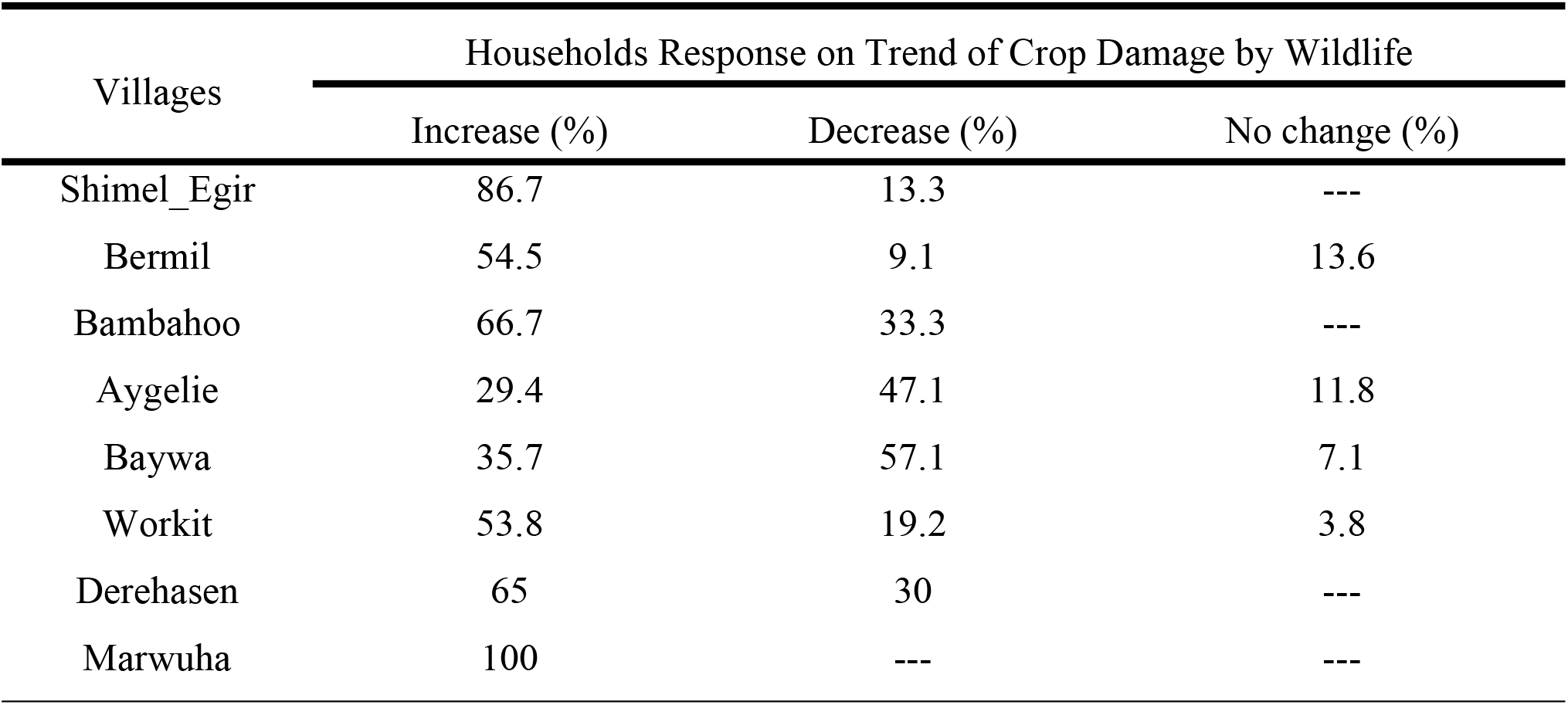
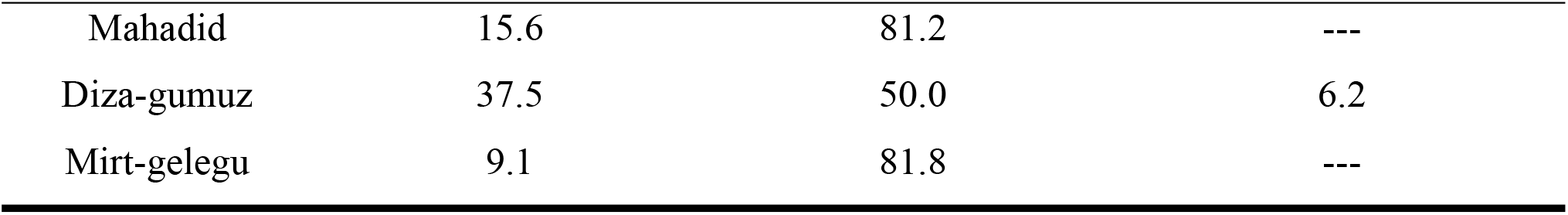
The perceived trends of wildlife caused crop damage across the villages

| Villages | Households Response on Trend of Crop Damage by Wildlife |  |  |
| --- | --- | --- | --- |
|  | Increase (%) | Decrease (%) | No change (%) |
| Shimel_Egir | 86.7 | 13.3 | --- |
| Bermil | 54.5 | 9.1 | 13.6 |
| Bambahoo | 66.7 | 33.3 | --- |
| Aygelie | 29.4 | 47.1 | 11.8 |
| Baywa | 35.7 | 57.1 | 7.1 |
| Workit | 53.8 | 19.2 | 3.8 |
| Derehasen | 65 | 30 | --- |
| Marwuha | 100 | --- | --- |
| Mahadid | 15.6 | 81.2 | --- |
| Diza-gumuz | 37.5 | 50.0 | 6.2 |
| Mirt-gelegu | 9.1 | 81.8 | --- |

### Effects of human-wildlife conflict on the local communities

Evidences from this study proved that human wildlife conflicts had various costs on communities living adjacent Alitash National park. The costs were diverse ranging from crop damage to lose of human lives. Crop damage was the leading cost in terms of prevalence and magnitude. Based on the respondents guess, a little more than one-sixth (18.7%) of the household was lost 10 to 25% of their crop fields to wild herbivores during the last 12 months prior to the survey period. A few (2%) of the surveyed households reported the highest (51 to 75 % crop field) level of crop damages in the study area during the same period (Table 5).

**Table 5:**
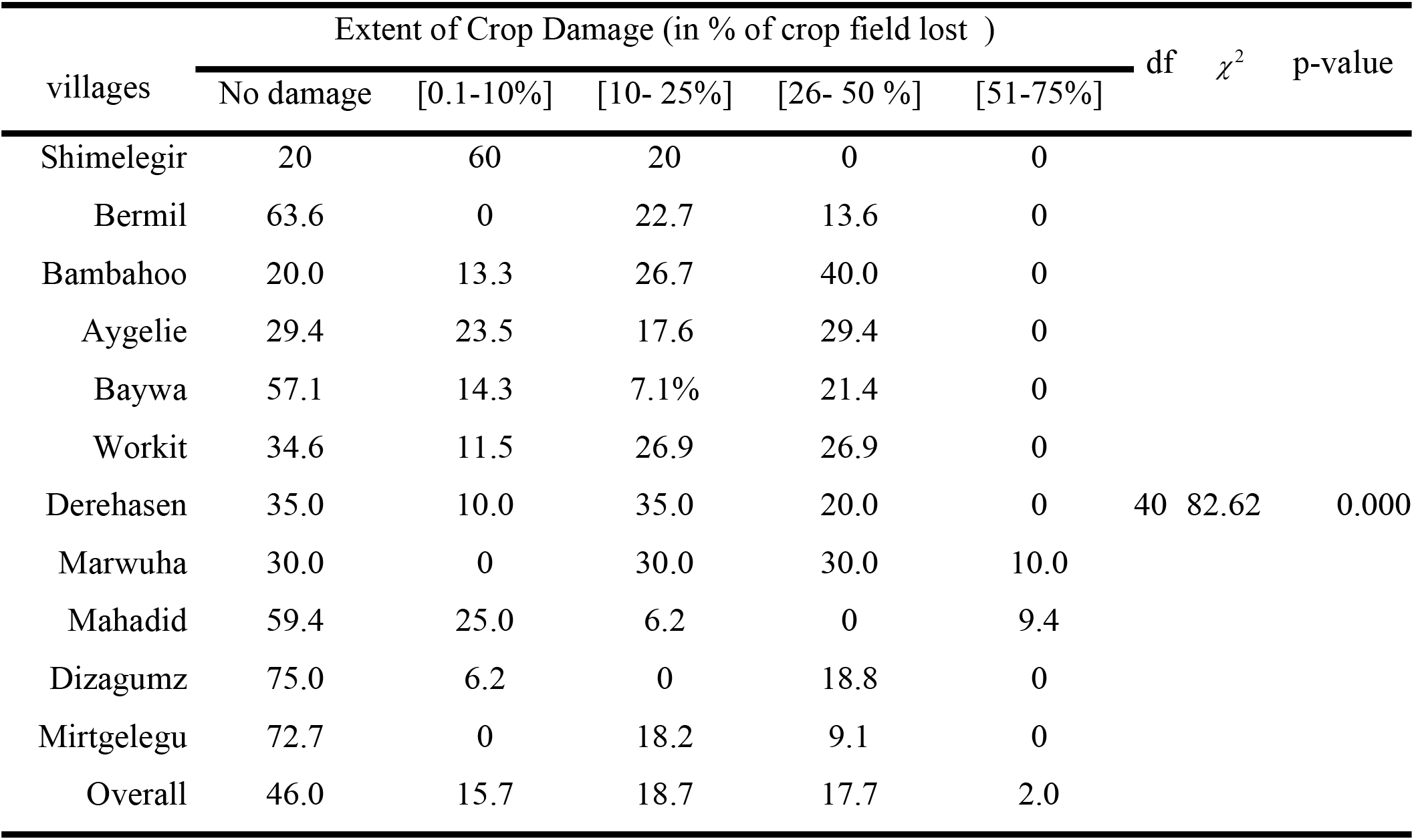
Proportion of respondents and their estimated crop loss

| villages | Extent of Crop Damage (in % of crop field lost ) | | | | | df | $\chi^2$ | p-value |
| --- | --- | --- | --- | --- | --- | --- | --- | --- |
|  | No damage | [0.1-10%] | [10- 25%] | [26- 50 %] | [51-75%] |  |  |  |
| Shimelegir | 20 | 60 | 20 | 0 | 0 | 40 | 82.62 | 0.000 |
| Bermil | 63.6 | 0 | 22.7 | 13.6 | 0 |  |  |  |
| Bambahoo | 20.0 | 13.3 | 26.7 | 40.0 | 0 |  |  |  |
| Aygelie | 29.4 | 23.5 | 17.6 | 29.4 | 0 |  |  |  |
| Baywa | 57.1 | 14.3 | 7.1% | 21.4 | 0 |  |  |  |
| Workit | 34.6 | 11.5 | 26.9 | 26.9 | 0 |  |  |  |
| Derehasen | 35.0 | 10.0 | 35.0 | 20.0 | 0 |  |  |  |
| Marwuha | 30.0 | 0 | 30.0 | 30.0 | 10.0 |  |  |  |
| Mahadid | 59.4 | 25.0 | 6.2 | 0 | 9.4 |  |  |  |
| Dizagumz | 75.0 | 6.2 | 0 | 18.8 | 0 |  |  |  |
| Mirtgelegu | 72.7 | 0 | 18.2 | 9.1 | 0 |  |  |  |
| Overall | 46.0 | 15.7 | 18.7 | 17.7 | 2.0 |  |  |  |

The extent of wildlife caused crop damages was significantly varied across the study villages (χ ^2^ = 82.62; df = 40; p = 0.000). The highest level crop damage (51 to 75% of crop fields) was only reported in Marwuha (10%) and Mahidid (9.4%) whereas the least (0.1-10% of crop fields) lose were reported by the majority (60%) of the respondents from Shimelegir. As compared with other villages, about a little more than one-third (35%) of the respondents from Derehasen encountered medium crop damage (10 to 25% of crop fields) within the last 12 months prior to the survey period.

The other cost of wildlife on the communities was livestock depredation. About 287.89 tropical livestock units (TLU) were lost due to wild predators during the last five years of period prior to the survey administration of this study. A little less than half (47.5%) of the surveyed households shared the loss which means 3.06 TLU on average. The overall mean loss livestock was 1.45TLU (Table 6).

**Table 6:** Livestock predated in the last five years prior to the study period

| Village | Sample | Livestock lost (in TLU) |  | St. deviation |
| --- | --- | --- | --- | --- |
|  |  | Total lose | Mean lose |  |
| Shimelegir | 15 | 43.02 | 2.87 | 3.02 |
| Bermil | 22 | 19.84 | 0.90 | 2.00 |
| Bambahoo | 15 | 47.18 | 3.15 | 2.53 |
| Aygelie | 17 | 42.99 | 2.53 | 2.84 |
| Baywa | 14 | 24.85 | 1.77 | 2.17 |
| Workit | 26 | 56.01 | 2.15 | 3.00 |
| Derehasen | 20 | 19.00 | 0.95 | 1.50 |
| Marwuha | 10 | 10.75 | 1.07 | 2.06 |
| Mahadid | 32 | 17.70 | 0.55 | 1.14 |
| Dizagumz | 16 | 3.30 | 0.21 | 0.53 |
| Mirtgelegu | 11 | 3.25 | 0.30 | 0.43 |
| Overall | 198 | 287.89 | 1.45 | 2.29 |

However, the distribution of livestock loss burden was varied among the study villages; the highest (56.01 TLU) lose was in Workit (56.01 TLU) followed by Bambahoo where the second peak (47.18 TLU) of loss was reported. Conversely, Bambahoo and Shimelegir were the first and the second with the mean loss of 3.15 and 2.87 TLU, respectively. According to the free list respondents, a total of 7 wild animals species were recorded as major predators of domestic animals in the study villages (Table 7).

**Table 7:**
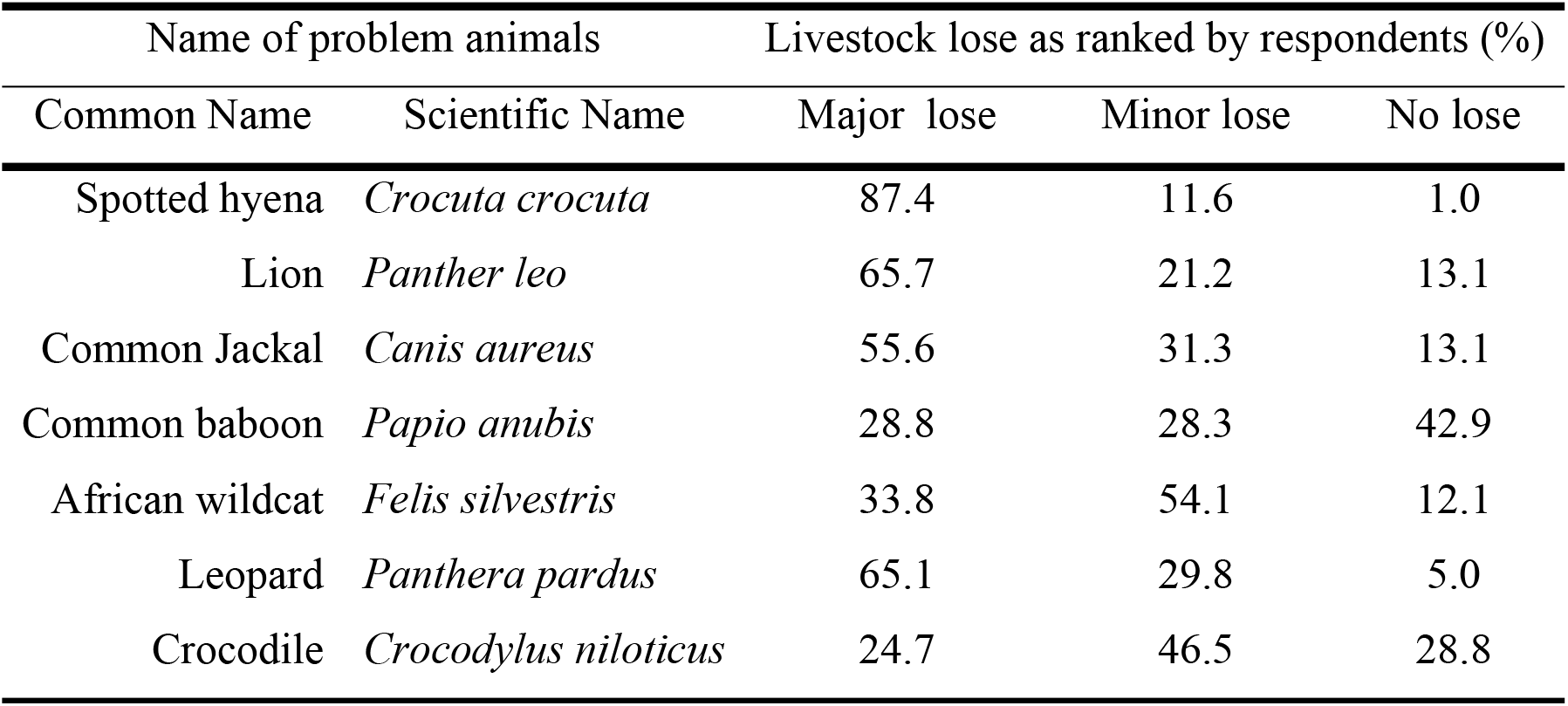
Major wild animals responsible for livestock predation

**Table 8:** Effect of wildlife caused damage on respondents’ perception towards wildlife

| Damage type | B | S.E. | Wald | df | Sig. | Exp(B) |
| --- | --- | --- | --- | --- | --- | --- |
| Livestock loss | -.250 | .081 | 9.607 | 1 | .002 | .779 |
| Crop Damage | -1.365 | .412 | 10.983 | 1 | .001 | .255 |
| Human Attack | -1.310 | .600 | 4.769 | 1 | .029 | .270 |

As confirmed by focus group discussants, attacks on people by wildlife were infrequent in the study area but serious and intolerable problem leading to strong public reactions. In fact, seven human attacks by lions, five by hyenas and six injuries by leopard were reported by respondents. According to the key informants, these attacks were happened when the victim cross the habitats of the animals, not in the victims residence.

Guarding crops or livestock was also reported as a cost of human–wildlife conflicts in the study communities. Respondents highlighted that how much time they waste for guarding instead of doing economic activities. This study also witnessed that guarding has a wide-range of intangible negative social and psychological impacts on human beings including fear, missing school and loss of sleep.

### Effects of HWC on wildlife Conservation

Wildlife conservation had been affected by human-wildlife conflict. A little less than one-third (32.3%) of the respondents had negative perceptions towards wildlife conservation efforts. A little more than two-third (79.7%) of them experienced wildlife caused damages. The key informants also confirmed that those encountered problem from wildlife never hesitate to take revenge either by directly killing the target problem animal or any individual of its species during the occasion or later on. In its worst case, the revenge could be deliberate fire setting to destruct their habitats.

To detect the major type of wildlife caused damages shaping perception of respondents towards wildlife conservation efforts, the study employed binary logistic regression model analysis. Indeed, three variables (livestock predation, crop damage and human attack) retained in the final model. The result of omnibus tests of the model coefficients showed that the probability of each step was <0.05 and the model coefficients were significant (*χ*^2^ = 46.845; df = 3; p = 0.000).. With respect to the goodness of fit of the model, the value of -2 log likelihood was 196.066 and the value of Nagelkerke *R* ^2^ was 0.298. The prediction accuracy of the model was 75.3%. Therefore, the model was reasonable to explain the significant variables (Table 9).

Livestock loss had a significant factor at 1 % significant level in influencing perception of local people towards wildlife and their conservation. Compared with the probability of respondents being negative to wildlife without encountering livestock loss, the probability of those loss livestock was higher. The odds ratio value suggests that when livestock losses raised by one unit (1TLU) the odds ratio became 0.779 times higher and hence the households are 0.779 times more likely to be negative for wildlife with other factors remains constant.

Likewise, wildlife caused crop damage had a significant and negative influence on perception of respondents towards wildlife conservation at 1% significance level. Comparing the probability of respondents to be negative for wildlife with and without facing wildlife caused crop damage; the probability of those encountered the damage became 0.255 times higher. The other major significant factor in shaping respondents’ perception towards wildlife conservation efforts was direct wildlife attack on human. Compared with probability of being negative to wildlife without attacks from wildlife, those encountered attack was 0.27times higher, if ceteris paribus.

## Discussions

The results suggest that human-wildlife conflicts have been a common phenomenon in and around Alitash National Park, which corroborates with previous findings (Abelieneh & Agitew, 2017; Berhanu & Teshome, 2018). Living around protected areas entails various types of conflicts with wildlife (Megaze et al., 2017) especially when the livelihoods of the communities depend mainly on agricultural activities like those living adjacent to Alitash National Park.

In the study area, both crop cultivation and livestock rearing play a vital role for the livelihood of the local people but highly prone to wildlife caused damages. Among the households surveyed, the majority were encountered crop and/or livestock damages due wildlife which had influence on the economic welfare of the local people. This finding concurs with previous studies in Ethiopia (Haylegebriel, 2016; Leta et al., 2016; Teshome & Girma, 2017) who reported crop damage and livestock depredation as a significant cost of farmers living close to wildlife habitats. Similarly Danson (et al., 2016), stated wildlife damages jeopardize socioeconomic wellbeing of people that depend on crop cultivation and livestock rearing for their livelihood.

Crop damage was most prevalent wildlife caused problem in the study area. This finding is in agreement with different studies in Ethiopia (Amare & Serekebirhan, 2019; Gebeyehu & Bekele 2009; Leta et al., 2016) which reported crop damage as the most serious problem in their respective study areas. As shown by Nyamwamu (2016) most farmers experience crop damage in Laikipia, Kenya where 10% of the farmers stop crop cultivation due to loss caused by wildlife. Since crop cultivation is the major livelihood strategy of the local people, even small damage on crop fields may highly affect the livelihood of those formers supporting their family by growing crops. Kabuusu (2018) reported that the destruction of crops as the worst form of human-wildlife conflict. Crop damages not only affects a farmer’s ability to feed his or her family, it is also reduces cash income and has repercussions for health, nutrition, education and ultimately, development. This is particularly heartbroken to the poorer farmers that are going to exploit wildlife and forest resources immediately after the damage happens to fulfill their household needs (Ogra, 2008).

The respondents of the study listed many wild animals responsible for crop damages in their locality. Among these common warthog (*Phacochoerus africanus*), African porcupines (*Hystrix cristata*), common baboon (*Papio cynocephalus Anubis*), vervet monkey (*Ceropithecus pygerythrus*) and patas monkey (*Erythrocebus patas*) were considered more problematic to the farming communities living around Alitash National Park. This matches with other findings in different parts of Ethiopia (Gebeyehu & Belkele, 2009; Fenta, 2014). Similar results were also observed on the study conducted in Southwestern Ethiopia which included common warthog (Phacochoerus africanus), grivet monkey (Cercopithecus aethiops) and crested porcupine (Hystrixcristata) in the list of most problematic wild animals causing crop damages (Gobosho et al., 2015).

Livestock depredation by wild predators was another form of human-wildlife conflict common in and around Alitash National Park. This fits with the fact that intensity of human–wildlife conflict is great around protected areas than elsewhere in any other places (Haylegebriel, 2016; Leta et al., 2016; Teshome & Girma, 2017). However, Amare & Serekebirhan (2019) in their study noted the least prevalence of livestock predation around Midre-Kebid Abo Monastry, Southwest Ethiopia. It can be inferred that keeping large number of livestock far from home may have high risk of losing livestock through predation in the study area.

Overall 287.89 tropical livestock units (TLU) were depredated by wild predators within the last five years prior to the survey administration of this study. High livestock lose were observed in Workit, Bambahoo and Shimelegir villages that are relatively far from the park. This finding contradicts with the finding of Acha & Temesgen (2015) which stated that high livestock losses observed in villages very close to Chebera-Churchura National Park. It can be inferred that there is difference in the methods of livestock keeping. Farmers in most of villages far from the park have a culture of keeping livestock away from their residences which give a great opportunity for wild predators to attack the livestock day and night. Livestock attacks not only associate with the remoteness but also the quality of the huts which are temporary and mobile with no well-built structures and fences to protect domestic animals against predators. Unlike those keeping livestock far away from their residence, those keeping livestock close to their dwellings have well-built permanent houses that can protect their livestock from occasionally roaming predators especially during the night. The closeness of the livestock houses to residences also gives a great chance of safely guarding livestock from predators.

The free list respondents showed that eight wild animals as major predators of domestic animals in the study villages. Spotted hyena (*Crocuta crocuta*), lion (*Panthera leo*) and common jackal (*Canis aureus*) were responsible for the most losses of livestock in the study area. Similar results were observed in and around Chebera-Churchura National Park (Acha & Temesgen, 2015).

The study also assessed the effect of human-conflict on perceptions of local people towards wildlife conservation. Majority of the respondents are positive for the conservation of wildlife in their locality. The result is similar to the finding of Megaze et al., (2017) who reported that most people around chebera churchura National Park were a positive towards the conservation area, despite the conflicts encountered with wildlife species of the area. This positive attitude among the respondent might be attributed to social and economic benefits they obtain from wildlife through illegal hunting and poaching practices which is impossible without the existence of wildlife in the respondent’s vicinity. As reported by Agitew& Abelieneh (2015), people living around Alitash National Park valued the park as source of livelihoods.

However, there were also responses indicating negative perceptions towards wildlife among the dwellers around the park. They consider wildlife unworthy for human being due to the damages incurred form problem animals. This is in agreement with Gebeyehu & Bekele (2009) who reported that the local people on Zegie Peninsula developed negative attitudes towards wild animals in response to the order of the problem they caused. Similarly, Eshete et al.(2018) in their study in south wollo, Ethiopia, stated a negative perception of people to the Ethiopian wolf due to its predation behavior on their sheep. Even those who faced low level of wildlife damages perceived wildlife as nuisance to their crops and enemy for their livestock. The result is also concur with Eniang et al.,(2011) who conveyed the role of wildlife caused damages to generate negative perceptions towards conservation.

In the study area, the negative perceptions towards wildlife were not just feeling but led people to kill wildlife and destroy their habitats. Particularly people who are dependent upon a single livelihood strategy tend to be more antagonistic towards wildlife, as the potential consequences of resource destruction are intensified by a lack of alternative assets or income strategies. This is in line with Biset et al.(2019), who reported retaliatory wildlife killing as a response to livestock depredation and crop damage.

## Conclusion and Recommendations

The current study examined effects of human wildlife conflicts on the livelihoods of the local people and wildlife conservation in and around Alitash National Park. Local Communities around Alitash National Park are negatively interacting with wildlife not only for their cultural practice but also due to their livelihoods. Crop cultivation and livestock rearing is playing a vital role for the livelihood of the local communities but also sources of conflicts between human and wildlife. Both people and wildlife incur tangible consequences from their conflicts. The presence of wildlife inflicts costs on local communities through crop damages, livestock predation and human attack. These affect not only the ability of a household to feed its family but also reduce cash income and have repercussions for health, nutrition, education and psychology.

Regardless of the multidimensional costs living alongside wildlife, the majority of the local people are positive towards wildlife conservation which provides a better chance of diminishing the negative interactions between people and wildlife through implementation of appropriate human-wildlife conflict prevention and mitigation strategies. However, the existing some negative perceptions among the local people have dire consequences on the effort of wildlife conservation in terms of retaliatory killings and ignorance of wildlife conservation.

As long as crop cultivation and livestock rearing remained means of livelihood with the usual practices, it is obvious that human-wildlife conflicts continue to negatively affect the welfare of people and wildlife. To this end, there is an urgent need for proper and effective land use planning whereby various land uses are specified and clearly demarcated to ensure settlements and other development projects far from the wildlife areas. Moreover, there is a need to protect rural livelihoods, reduce their vulnerability, and counterbalance losses with benefits from wildlife and nature tourism. Alternative compensation system should be applied to those lost their crops and/or livestock by the activities of wildlife unless retaliatory killings will continue and results wildlife decline and species extinction.

## Acknowledgement

We are grateful to University of Gondar for providing the fund and facilities. We thank staffs of Alitash National Park for their overall assistance and facilitation throughout the research period. We would also like to acknowledge all the study participants for their patience and time they sacrificed and all those who helped us during the study.

## Notes

### Competing Interest Statement

The authors have declared no competing interest.

